# Resurrecting essential amino acid biosynthesis in a mammalian cell

**DOI:** 10.1101/2021.08.03.454854

**Authors:** Julie Trolle, Ross M. McBee, Andrew Kaufman, Sudarshan Pinglay, Henri Berger, Sergei German, Liyuan Liu, Michael J. Shen, Xinyi Guo, J. Andrew Martin, Michael Pacold, Drew R. Jones, Jef D. Boeke, Harris H. Wang

## Abstract

Major genomic deletions in independent eukaryotic lineages have led to repeated ancestral loss of biosynthesis pathways for nine of the twenty canonical amino acids^1^. While the evolutionary forces driving these polyphyletic deletion events are not well understood, the consequence is that extant metazoans are unable to produce nine essential amino acids (EAAs). Previous studies have highlighted that EAA biosynthesis tends to be more energetically costly^2,3^, raising the possibility that these pathways were lost from organisms with access to abundant EAAs in the environment^4,5^. It is unclear whether present-day metazoans can reaccept these pathways to resurrect biosynthetic capabilities that were lost long ago or whether evolution has rendered EAA pathways incompatible with metazoan metabolism. Here, we report progress on a large-scale synthetic genomics effort to reestablish EAA biosynthetic functionality in a mammalian cell. We designed codon-optimized biosynthesis pathways based on genes mined from *Escherichia coli*. These pathways were *de novo* synthesized in 3 kilobase chunks, assembled *in yeasto* and genomically integrated into a Chinese Hamster Ovary (CHO) cell line. One synthetic pathway produced valine at a sufficient level for cell viability and proliferation, and thus represents a successful example of metazoan EAA biosynthesis restoration. This prototrophic CHO line grows in valine-free medium, and metabolomics using labeled precursors verified *de novo* biosynthesis of valine. RNA-seq profiling of the valine prototrophic CHO line showed that the synthetic pathway minimally disrupted the cellular transcriptome. Furthermore, valine prototrophic cells exhibited transcriptional signatures associated with rescue from nutritional starvation. This work demonstrates that mammalian metabolism is amenable to restoration of ancient core pathways, thus paving a path for genome-scale efforts to synthetically restore metabolic functions to the metazoan lineage.

---

Whole genome sequencing across the tree of life has revealed the surprising observation that nine amino acid (AA) biosynthesis pathways are missing from the metazoan lineage^1^. Furthermore, these losses appear to have occurred multiple times during eukaryotic evolution, including in some microbial lineages (**Fig 1A**)^1,4^. Branching from core metabolism, the nine EAA biosynthesis pathways missing from metazoans involve over forty genes (**Fig 1B, Table ED1-ED3**), widely found in bacteria, fungi and plants^4^. While the absence of essential metabolic pathways is observed in certain bacteria^6^, which possess short generation times and high genomic flexibility to adapt to rapidly changing environments, the forces driving the loss of multiple EAA biosynthetic pathways in multicellular eukaryotes remain a great mystery. Their partial reacquisition through horizontal gene transfer in certain rare insect lineages with extremely simple nutrient sources, such as sap or blood, enables them to host genome-reduced intracellular bacteria that provide other missing metabolites missing from these limited diets, and provides the “exception that proves the rule”^7^. Recent efforts in genome-scale synthesis^8-10^ and genome-writing^11^ have highlighted our increasing capacity to construct synthetic genomes with novel properties, thus providing a route to not only examine these interesting evolutionary questions but also yield new capacities of bioindustrial utility ^12-14^.

**Figure 1.**
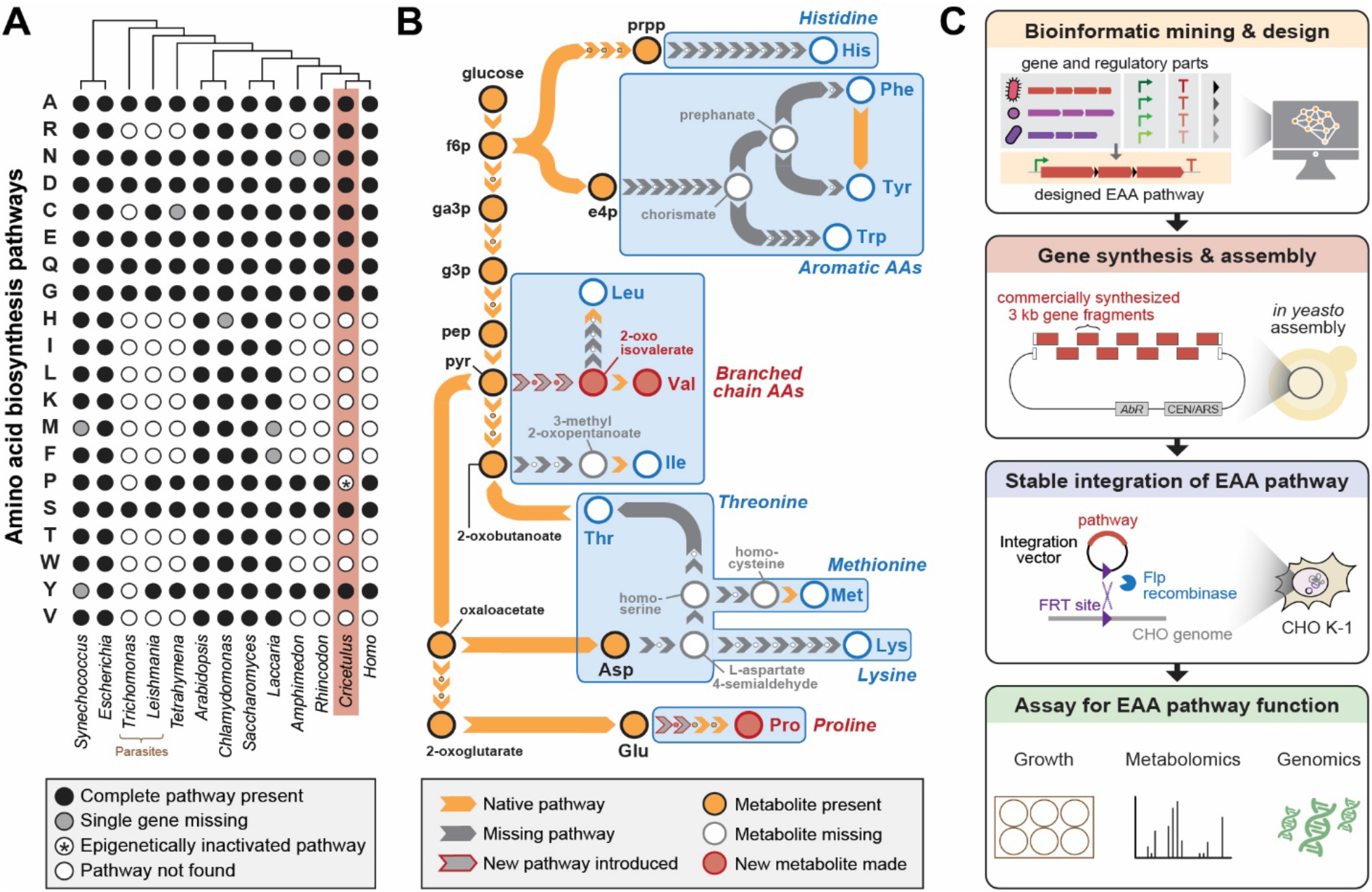
Engineering Essential Amino Acid (EAA) biosynthesis in metazoan cells. **(A)** Presence of amino acid biosynthesis pathways across representative diverse organisms on Earth. **(B)** Schematic of EAA biosynthesis pathway steps that require engineering in mammalian cells to enable complete amino acid prototrophy if imported from *E. coli*. Proline and Valine pathways shown in this work are highlighted in red. **(C)** Workflow diagram of a synthetic genomics approach pathway design, construction, integration and testing towards mammalian EAA restoration.

We sought to explore the possibility of generating prototrophic mammalian cells capable of complete biosynthesis of EAAs using a synthetic genomics approach (**Fig 1C**). The Chinese Hamster Ovary (CHO) K1 cell line was chosen as a model system due to its fast generation time, amenability to genetic manipulations, availability of a whole genome sequence, and established industrial relevance for producing biologics^15^. EAA biosynthesis genes from the best characterized model organisms were considered during pathway design while optimizing for the fewest number of enzymes needed for a given EAA pathway. To avoid using multiple promoters, we introduced ribosome-skipping 2A sequences^16^ between biosynthetic genes to allow for protein translation of separate enzymes from a single transcriptional unit. The EAA pathway and an additional EGFP reporter were placed in a vector that could be integrated as a single copy into the CHO genome at a designated landing pad using the FLP-In system^17^. The entire pathway was synthesized *de novo* by commercial gene synthesis in 3 kilobase fragments and assembled in *Saccharomyces cerevisiae* via homologous recombination of 80 basepair overlaps. Subsequent antibiotic selection of cells transfected with the vector resulted in a stable cell line containing the integrated EAA pathway. Finally, we performed a variety of phenotypic, metabolomic, and transcriptomic characterizations on the modified cell line to verify activity of the EAA biosynthesis pathway.

We first confirmed that the CHO cell line was auxotrophic for each of the 9 EAAs. As expected, CHO-K1 did not grow in “dropout” F-12K medium lacking each of the 9 EAAs and supplemented with dialyzed FBS (**Fig S1**). We noted that in this cell line, canonically non-essential amino acids tyrosine and proline also exhibited EAA-like properties in dropout media. Insufficient concentrations of phenylalanine in F-12K media or low expression of endogenous phenylalanine-4-hydroxylase that converts phenylalanine to tyrosine could underlie tyrosine limitation. Proline auxotrophy in CHO-K1 results from epigenetic silencing of the gene encoding Δ1-pyrroline-5-carboxylate synthetase (P5CS) in the proline pathway^18^. We therefore used proline as a test case for our synthetic genomics pipeline. We tested the P5CS-equivalent proline biosynthesis enzyme found in *E. coli*, encoded by two separate genes, *proA* and *proB* (**Fig ED2A**). A vector (pPro) carrying codon-optimized *proA* and *proB* separated by a P2A sequence was synthesized and integrated into CHO-K1 (**Fig ED2B**). CHO cells with the stably integrated pPro proline pathway showed robust growth in proline-free medium (**Fig ED2C-D**), thus validating a pipeline for designing and generating specific AA prototrophic cells.

To demonstrate restoration of EAA pathways lost from the metazoan lineage more than 650-850 million years ago^19^, we built a 6-gene construct (pMTIV) to test the simultaneous rescue of methionine, threonine, isoleucine and valine auxotrophies. These EAAs were chosen because their biosynthesis pathways were missing the fewest number of genes: methionine and threonine production require two genes while valine and isoleucine require four genes total (**Fig ED3**). To biosynthesize methionine, we chose the *E. coli metC* gene, which converts cystathionine to homocysteine, a missing step in CHO-K1 cells in a potential serine to methionine biosynthetic pathway. Threonine production was tested using *E. coli* glycine hydroxymethyltransferase *ltaE*, which converts glycine and acetaldehyde into threonine. For branched chain amino acids (BCAAs) valine and isoleucine, three additional biosynthetic enzymes and one regulatory subunit are needed in theory to convert pyruvate and 2-oxobutanoate into valine and isoleucine, respectively. In the case of valine, pyruvate is converted to 2-acetolactate, then to 2,3-dihydroxy-isovalerate, then to 2-oxoisovalerate and finally to valine. For isoleucine, 2-oxobutanoate is converted to 2-aceto-2-hydroxybutanoate, then to 2,3-dihydroxy-3-methylpentanoate, then to 3-methyl-2-oxopentanoate, and finally to isoleucine. The final steps in both BCAAs can be performed by native CHO catabolic enzymes Bcat1 and Bcat2^18^. In *E. coli*, the first three steps in the pathway are embodied in four genes that encode an acetolactate synthase split into catalytic and regulatory subunits (*ilvB/N*), a ketol-acid reductoisomerase (*ilvC*), and a dihydroxy-acid dehydratase (*ilvD*) (**Fig 2A**)^20^. The final pMTIV construct comprises *metC, itaE, ilvB, ilvN, ilvC* and *ilvD*, organized as a single open reading frame (ORF) with a 2A sequence variant in between each protein coding region (**Fig 2B**), driven by a strong SFFV viral promoter.

**Figure 2.**
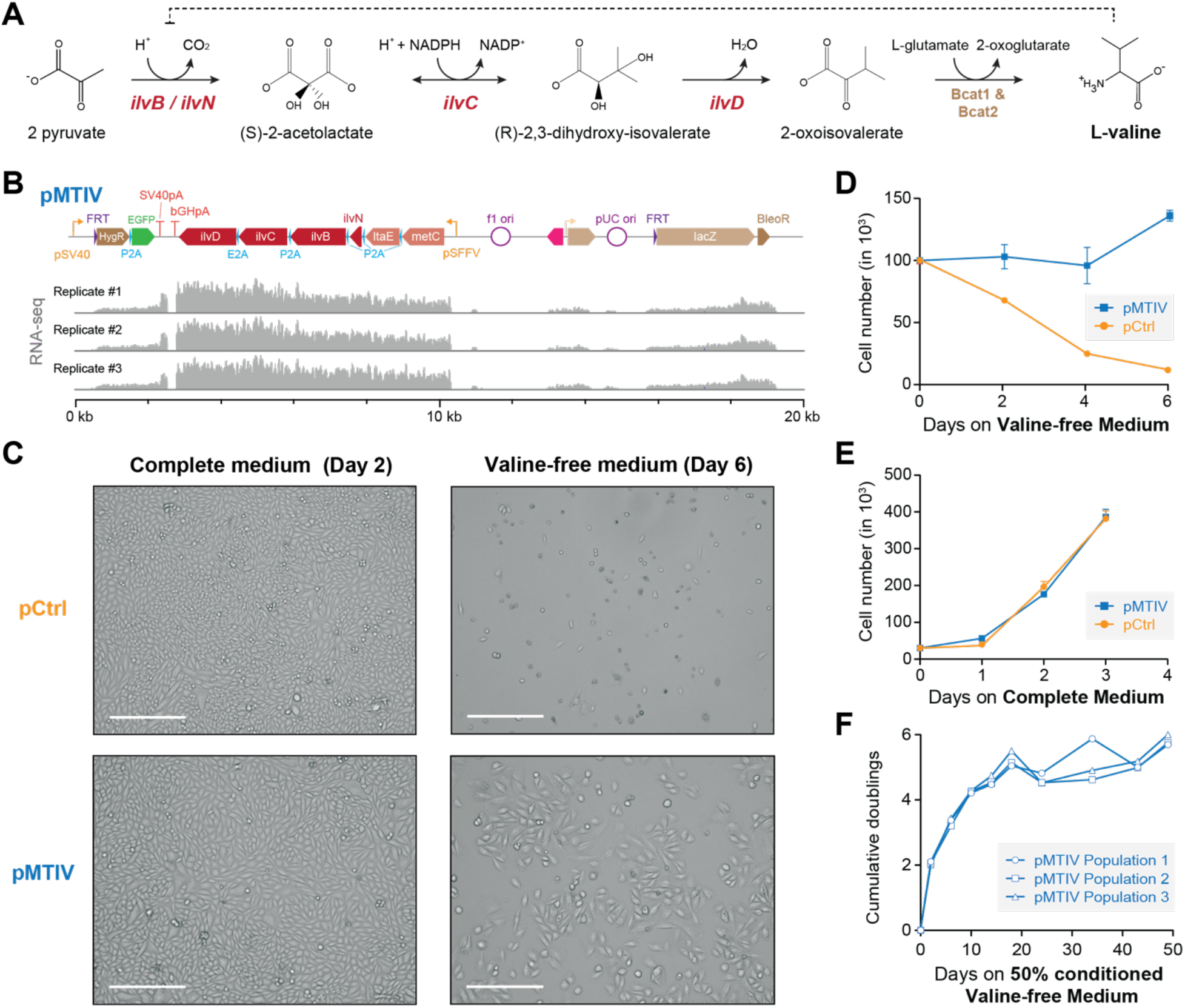
Restoration of a Valine biosynthesis pathway in CHO-K1 cells. **(A)** Three enzymatic steps encoded by *E. coli* genes *ilvN, ilvB, ilvC*, and *ilvD* are required for valine biosynthesis in CHO-K1 cells. **(B)** Schematic of pMTIV construct after genomic integration and RNA-seq read coverage showing successful incorporation and active transcription. **(C)** Microscopy images of CHO-K1 cells with integrated pCtrl or pMTIV constructs in Complete Medium after 2 days or valine-free medium after 6 days. Scale bar = 300 um. **(D)** Growth curve of CHO-K1 cells with pCtrl or pMTIV in valine-free medium. Day 0 indicates number of seeded cells. Triplicate population were plated for each timepoint in each of three 24-well wells, error bars show the standard deviation or replicates. **(E)** Growth curve of CHO-K1 cells with pCtrl or pMTIV in complete medium. Day 0 indicates number of seeded cells. Triplicate populations were plated for each timepoint in each of three 24-well wells. Error bars show the standard deviation of replicates **(F)** CHO-K1 with pMTIV cultured over 9 passages in 50% conditioned valine-free medium.

To test the biosynthetic capacity of pMTIV, we first introduced the construct into CHO cells. Flp-In integration was used to stably insert either pMTIV, or a control vector (pCtrl) into the CHO genome. Successful generation of each cell line was confirmed by PCR amplification of junction regions formed during vector integration (**Fig ED4A-B**). RNA-seq of cells containing the pMTIV construct confirmed transcription of the entire ORF (**Fig 2C**). Western blotting of pMTIV cells using antibodies against the P2A peptide yielded bands at the expected masses of P2A-tagged proteins, confirming the production of separate distinct enzymes (**Fig ED4C**).

In methionine-free, threonine-free, or isoleucine-free medium, cells containing the pMTIV construct did not show viability over seven days, similar to cells containing the pCtrl control vector **Fig ED5**). In striking contrast, however, cells containing the integrated pMTIV showed relatively healthy cell morphology and viability in valine-free medium (**Fig 2D**), whereas cells containing pCtrl exhibited substantial loss of viability over six days. In complete medium, cells carrying the integrated pMTIV construct showed no growth defects compared to control cells (**Fig 2E**). In valine-free medium, pMTIV cells showed a 32% increase in cell number over 6 days compared to an 88% decrease in cell number in pCtrl cells (**Fig 2F**). When cultured in valine-free medium over multiple passages with medium changes every two days, pMTIV cell proliferation was substantially reduced by the 3^rd^ passage. We hypothesized that frequent passaging might over-dilute the medium and prevent sufficient accumulation of biosynthesized valine necessary for continued proliferation. We thus deployed a “conditioned-medium” regimen whereby 50% of the medium was freshly prepared valine-free medium and 50% was “conditioned” valine-free medium in which pMTIV cells had previously been cultured over 2 days (see Methods). Using this regimen, we were able to culture pMTIV cells for 9 passages without addition of exogenous valine, during which time they exhibited an average doubling time of 8.5 days. However, the doubling time varied across the 49 days of experimentation with cells exhibiting a mean doubling time of 5.3 days in the first 24 days and 21.0 days in the last 25 days. The increase in doubling time seen in later passages may be the result of detrimental effects from culturing cells longer-term in partially recycled and dialyzed FBS or may result from variation in the cell number to medium volume ratio, which trended downwards in later passaging events as cell growth slowed. Despite the slowed growth seen in later passages, cells exhibited healthy morphology and continued to proliferate at day 49, suggesting that the cells could have been passaged even further. To verify that the putative valine rescue effect was due to the valine biosynthesis genes present in pMTIV specifically, we constructed and tested a second EAA pathway vector pIV that only contained the four genes *ilvNBCD*. The pIV construct similarly supported cell growth in valine-free medium, and exhibited similar growth dynamics to the pMTIV construct in complete medium (**Fig ED5, Fig ED6**).

To confirm endogenous biosynthesis of valine, we cultured pCtrl and pMTIV cells in RPMI medium containing ^13^C_6_-glucose in the place of its ^12^C equivalent together with ^13^C_3_-pyruvate spiked in at 2 mM over 3 passages (**Fig ED7A**). High-resolution MS1 of MTIV cell lysates revealed a peak at 123.1032 *m*/*z*, the expected *m*/*z* for ^13^C_5_-valine (**Fig 3A**). This detected peak was subject to MS2 alongside a ^12^C-valine control peak and a ^13^C_5_/^15^N-valine peak, which was spiked into all samples to serve as an internal standard. The resulting fragmentation patterns for each peak (**Fig 3B**) matched the theoretical expectations for each isotopic version of valine (**Fig ED7B**). An extracted ion chromatogram further revealed a peak in the pMTIV valine-free medium metabolite extraction, which corresponded to a peak in the spiked-in positive control ^13^C_5_/^15^N-valine, whereas no equivalent peak was seen among metabolites extracted from pCtrl cells (**Fig ED7C**). Taken together, this demonstrates that pMTIV cells are biosynthesizing valine from core metabolites glucose and pyruvate, thereby representing successful metazoan biosynthesis of valine. Over the course of 3 passages in heavy valine-free medium, the non-essential amino acid alanine, which is absent from RPMI medium and synthesized from pyruvate, was found to be 86.1% ^13^C-labeled in pMTIV cell lysates. Assuming similar turnover rates for alanine and valine within the CHO proteome, we expected to see similar percentages of ^13^C-labeled valine. However, just 32.2% of valine in pMTIV cell lysates was ^13^C-labeled (**Fig ED7D-E**). For pMTIV cells cultured in heavy complete medium, just 6.4% of valine in cell lysates was ^13^C-labeled. Together with the observed slow proliferation of pMTIV cells in valine-free medium, our data suggests that valine complementation is sufficient but perhaps sub-optimal for cell growth.

**Figure 3.**
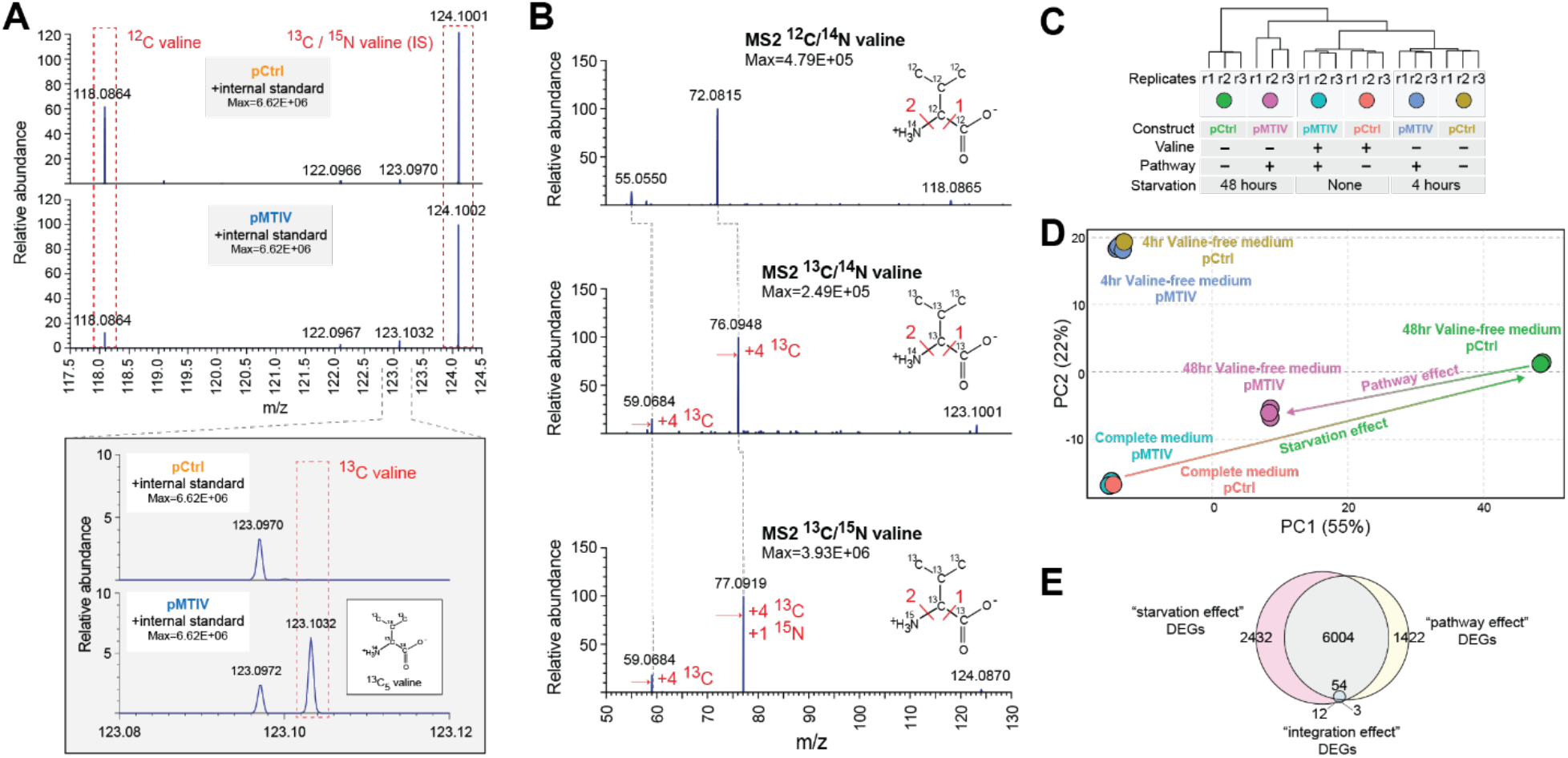
Heavy-carbon validation of endogenous valine production and transcriptomic signatures associated with rescue of nutritional starvation. **(A)** Mass spectra showing an MS1 peak corresponding to the expected *m*/*z* for ^13^C-valine when pMTIV cells were grown on valine-free medium supplemented with ^13^C-glucose and ^13^C-pyruvate, indicating autogenous production of intracellular valine. **(B)** MS2 performed on peaks extracted from pMTIV sample, which corresponded to *m*/*z* values expected for ^12^C-valine, ^13^C-valine and internal standard ^13^C/^15^N-valine. MS2 fragmentation patterns for each of these metabolites matched expectations. **(C)** RNAseq dendrogram of cells with and without the pMTIV pathway grown on complete medium or starved of valine for 4 or 48 hours. **(D)** PCA space depiction of cells with or without the pMTIV pathway grown on complete medium, or starved of valine for 4 or 48 hours. **(E)** Overlap between Differently Expressed Genes (DEGs) comparing cells without the pathway cultured in valine-free medium to cells cultured in complete medium (the “starvation effect”), cells with and without the pMTIV pathway after 48 hours starvation on valine-free medium (the “pathway effect”), and cells with and without the pathway grown on complete medium (the “integration effect”).

We performed RNA-seq to profile the transcriptional responses of cells containing pMTIV or pCtrl in complete (harvested at 0 h) and valine-free medium (harvested at 4 and 48 h, respectively) (**Fig 3C, Fig ED8A**). The transcriptional impact of pathway integration is modest (**Fig 3D**). Only 51 transcripts were differentially expressed between pCtrl and pMTIV cells grown in complete medium, and the fold changes between conditions were small (**Fig 3E, Fig ED8B**). While some gene ontology (GO) functional categories were enriched (**Fig ED8C**), they did not suggest dramatic cellular stress. Rather, these transcriptional changes may reflect cellular response to BCAA dysregulation due to alterations in valine concentrations^21^, or they may result from cryptic effects of bacterial genes placed in a heterologous mammalian cellular context. In contrast, comparison of 48 h valine-starved pCtrl and pMTIV cells yielded ∼7,500 differentially expressed genes. Transcriptomes of pMTIV cells in valine-free medium more closely resembled cells grown on complete medium than did pCtrl cells in valine-free medium (**Fig 3D, Fig ED9A)**. Differentially expressed genes between pCtrl and pMTIV cells showed enrichment for hundreds of GO categories, including clear signatures of cellular stress such as autophagy, changes to endoplasmic reticulum trafficking, and ribosome regulation (**Fig ED9B**). Most of the differentially regulated genes between pCtrl cells in complete medium, and those same cells starved of valine for 48 hours were also differentially expressed when comparing pCtrl and pMTIV cells in valine-free medium (**Fig 3E**), supporting the hypothesis that most of the observed transcriptional changes represent broad but partial rescue of the cellular response to starvation.

In this work, we demonstrated the successful restoration of an EAA biosynthetic pathway in a metazoan cell. Our results indicate that contemporary metazoan biochemistry can support complete biosynthesis of valine, despite millions of years of evolution from its initial loss from the ancestral lineage. Interestingly, independent evidence for BCAA biosynthesis has also been obtained for sap-feeding whitefly bacteriocytes that host bacterial endosymbionts; metabolite sharing between these cells is predicted to lead to biosynthesis of BCAAs that are limiting in their restricted diet. The malleability of mammalian metabolism to accept heterologous core pathways opens up the possibility of animals with designer metabolisms and enhanced capacities to thrive under environmental stress and nutritional starvation^22^. Yet, our failure to functionalize designed methionine, threonine and isoleucine pathways highlights outstanding challenges and future directions. Other pathway components or alternative selections may be needed for different EAAs^23^. A general lack of predictability and a dearth of well-characterized and controllable genetic “parts” with high dynamic range continue to hamper efforts in genome-scale mammalian engineering^24-26^. Studies to reincorporate EAAs into the core mammalian metabolism could provide greater understanding of nutrient-starvation in different physiological contexts including the tumor microenvironment^27^, help answer deep evolutionary questions regarding the formation of the metazoan lineage^28^, and lead to new model systems or even therapeutics to address metabolic syndrome, Maple Syrup Urine Disease^29^ and Phenylketonuria^30^ all of which involve amino acid biosynthetic dysfunction^31,32^. Emerging synthetic genomic efforts to build a prototrophic mammal may require reactivation of many more genes (**Table ED1-ED3**), iterations of the design, build, test (DBT) cycle, and a larger coordinated research effort to ultimately bring such a project to fruition.

## METHODS

### Pathway completeness analysis

For pathway completeness analysis, the EC numbers of each enzyme in each amino acid biosynthesis pathway (excluding pathways annotated as only occurring in prokaryotes) were collected from the MetaCyc database (**Table ED4**). Variant biosynthetic routes to the same amino acid were considered as separate pathways, generating distinct EC number lists. The resulting per-pathway EC number lists were checked against the KEGG, Entrez Gene, Entrez Nucleotide, and Uniprot databases using their respective web APIs for each listed organism. If the combination of all databases contained at least one complete EC numbers list, corresponding to an end-to-end complete biosynthetic pathway, the organism was considered “complete” for that essential amino acid.

### Cell lines and media

CHO Flp-In™ cells (ThermoFisher, R75807) were used in all experiments. For growth assays involving amino acid dropout formulations, medium was prepared from an amino acid-free Ham’s F-12 (Kaighn’s) powder base (US Biological, N8545), and custom combinations of amino acids were added back in as needed to match the standard amino acid concentrations for Ham’s F-12 (Kaighn’s) medium or as specified. Custom amino acid dropout medium was adjusted to a pH of 7.3, sterile filtered, and supplemented with 10% dialyzed Fetal Bovine Serum and Penicillin-Streptomycin (100U/mL) prior to use. For metabolomics experiments, medium was prepared from an amino acid-free and glucose-free RPMI 1640 powder base (US Biological, R9010-01), and custom combinations of amino acids and isotopically heavy glucose and sodium pyruvate were added in to match the standard amino acid concentrations for RPMI 1640 or as specified. pH was adjusted to 7.3, sterile filtered and supplemented with 10% dialyzed Fetal Bovine Serum and Penicillin-Streptomycin (100U/mL) prior to use.

### Cell counting and quantification

For amino acid dropout curves, cells were seeded at 1×10^4^ into 6-well plates into F12-K media with lowered amino acid concentrations relative to typical F12-K media and then allowed to grow for five days. Media was then aspirated off and replaced with PBS with Hoechst 33342 live nuclear stain for automated imaging and counting using a DAPI filter set on an Eclipse Ti2 automated inverted microscope. To count, an automated microscopy routine was used to image 5 random locations within each well at 10x magnification, and then the cells present in imaged frames counted using automatic cell segregation and counting software. Given differences in cell response to starvation, segregation and counting parameters were tuned each experiment, but kept constant between starvation conditions and cells with and without the pathway. For synthetic prototroph pathway tests, raw cell counts were performed using the Countess II Automated Cell Counter (ThermoFisher, A27977) in accordance with the manufacturer’s protocol, or using the Scepter 2.0 Handheld Automated Cell Counter (Milipore Sigma, C85360) in accordance with the manufacturer’s protocol. Where indicated, relative cell quantification was measured using PrestoBlue™ Cell Viability Reagent (ThermoFisher, A13261) in accordance with the manufacturer’s protocol.

### Culturing synthetic prototrophic cells without exogenous supply of valine

For long-term culture of synthetic prototrophic cells, cells were cultured in 50% conditioned valine-free F12-K medium. Conditioned medium was generated by seeding 1×10^6^ pMTIV cells into 10mL complete F12-K medium on 10cm plates and replacing the medium with 10mL freshly prepared valine-free F12-K medium the next day following a PBS wash step. Cells conditioned the medium for 2 days at which point the medium was collected, centrifuged at 300xg for 3 mins to remove potential cell debris, and collected in 150mL vats to reduce batch-to-batch variation. This 100% conditioned medium was subsequently mixed in a 1:1 ratio with freshly prepared, unconditioned valine-free medium to generate so-called 50% conditioned valine-free medium, which was used throughout the long-term culturing process of synthetic prototrophic cells without exogenous supply of valine, For long-term culturing, 1×10^5^ pMTIV cells were seeded in triplicate populations into complete medium in 6-well plates, (day -1) which was replaced with 50% conditioned valine-free medium the next day (day 0) following a PBS wash step. Cells were counted at each passaging event and split at a 1:4, 1:2 or 3:4 proportion such that approximately 2×10^6
^ cells were seeded at each passaging event as best possible.

### DNA assembly, recovery and amplification

Integrated constructs were synthesized *de novo* in 3kb DNA segments with each segment overlapping neighboring segments by 80. Assembly was conducted *in yeasto* by co-transformation of segments into *Saccharomyces cerevisiae*. After 2 days of selection at 30C on Sc-URA, individual colonies were picked and cultured overnight. 1.5mL of each resulting yeast culture was resuspended in 250ul of P1 resuspension buffer (Qiagen, 19051) containing RNase. Glass beads were added to each resuspension and the mixture was vortexed for 10 mins to mechanically shear the cells. Next, cells were subject to alkaline lysis by adding 250ul of P2 lysis buffer (Qiagen, 19052) for 5 mins and then neutralized by addition of Qiagen N3 neutralization buffer (Qiagen, 19053). Subsequently, cell debris was spun down and plasmid DNA was collected using the Zymo Zyppy plasmid preparation kit (Zymo Research, D4036) according to the manufacturer’s instructions. Plasmid DNA was eluted in 30ul of Zyppy Elution buffer of which 10ul would be transformed into 100ul of *E. coli* for plasmid amplification.

### Protein extraction and western blot

Cell were lysed in SKL Triton lysis buffer (50 mM Hepes pH7.5, 150 mM NaCl, 1 mM EDTA, 1 mM EGTA, 10% glycerol, 1% Triton X-100, 25 mM NaF, 10 μM ZnCl_2_) supplemented with protease inhibitor (Sigma 11873580001). NuPAGE™ LDS sample buffer (ThermoFisher, NP0007) supplemented with 1.43 M β-mercaptoethanol was added to samples prior to heating at 70C for 10 mins. Gel electrophoresis was performed using 4-12% Bis-Tris gels (ThermoFisher, NP0326BOX) and run in NuPAGE™ MOPS running buffer (ThermoFisher, NP0001). Proteins were then transferred onto a PVDF membrane (Milipore Sigma, IPFL00010) using the Biorad Trans-Blot Turbo system in accordance with the manufacturer’s instructions. The transfer membrane was blocked in Odyssey blocking buffer (LI-COR, 927-40000) for 1 h at room temperature prior to incubation in primary antibody (Novus Biologicals, NBP2-59627 [1:1000 dilution]; Cell Signaling Technology, 2148 [1:1000 dilution]) solubilized in a 1:1 mixture of Odyssey blocking buffer and TBS-T buffer (50 mM Tris Base, 154 mM NaCl, 0.1% Tween20) overnight at 4C. Secondary antibodies (LI-COR, 926-32210 [1:20,000 dilution]; LI-COR, 926-68071 [1:20,000 dilution]),were also solubilized in Odyssey blocking buffer / TBS-T buffer. The membrane was incubated in the secondary antibody solution for 1.5 h at room temperature.

### Metabolomics

Cells were cultured in IH medium over 3 passages prior to cell harvest. Cell pellets were generated by trypsinization, followed by low speed centrifugation, and the pellet was frozen at - 80°C until further processing. A metabolite extraction was carried out on each sample with an extraction ratio of 1e6 cells per mL (80% methanol containing internal standards, 500 nM), according to a previously described method^33^. The LC column was a Millipore™ ZIC-pHILIC (2.1 x150 mm, 5 μm) coupled to a Dionex Ultimate 3000™ system and the column oven temperature was set to 25°C for the gradient elution. A flow rate of 100 μL/min was used with the following buffers; A) 10 mM ammonium carbonate in water, pH 9.0, and B) neat acetonitrile. The gradient profile was as follows; 80-20%B (0-30 min), 20-80%B (30-31 min), 80-80%B (31-42 min). Injection volume was set to 1 μL for all analyses (42 min total run time per injection). MS analyses were carried out by coupling the LC system to a Thermo Q Exactive HF™ mass spectrometer operating in heated electrospray ionization mode (HESI). Method duration was 30 min with a polarity switching data-dependent Top 3 method for both positive and negative modes, and targeted MS2 scans for the monoisotopic, U-13C, and U-13C/U-15N valine *m*/*z* values. Spray voltage for both positive and negative modes was 3.5kV and capillary temperature was set to 320°C with a sheath gas rate of 35, aux gas of 10, and max spray current of 100 μA. The full MS scan for both polarities utilized 120,000 resolution with an AGC target of 3e6 and a maximum IT of 100 ms, and the scan range was from 67-1000 *m*/*z*. Tandem MS spectra for both positive and negative mode used a resolution of 15,000, AGC target of 1e5, maximum IT of 50 ms, isolation window of 0.4 m/z, isolation offset of 0.1 m/z, fixed first mass of 50 m/z, and 3-way multiplexed normalized collision energies (nCE) of 10, 35, 80. The minimum AGC target was 1e4 with an intensity threshold of 2e5. All data were acquired in profile mode. All valine data were processed using Thermo XCalibur Qualbrowser for manual inspection and annotation of the resulting spectra and peak heights referring to authentic valine standards and labeled internal standards as described.

### RNA Seq

RNA was extracted from cells using the Qiagen RNeasy Kit (Qiagen, 74104) according to the manufacturer’s protocol. QIAshredder homogenizer columns were used to disrupt the cell lysates (Qiagen, 79654). mRNA was purified using the NEBNext poly(A) mRNA Magnetic Isolation module (New England Biolabs, E7490) in accordance with the manufacturer’s protocol. Libraries were prepared using the NEBNext Ultra RNA Library Prep Kit for Illumina (New England Biolabs, E7770), and sequenced on a NextSeq 550 single-end 75 cycles high output with v2.5 chemistry. Reads were adapter and quality trimmed with fastP using default parameters and psuedoaligned to the GCF_003668045.1_CriGri-PICR Chinese hamster genome assembly using kallisto. Differential gene enrichment analysis was performed with in R with DESeq2 and GO enrichment performed and visualized with clusterProfiler against the org.Mm.eg.db database, with further visualization with the pathview, GoSemSim, eulerr packages.

## Acknowledgements

We would like to thank the members of the Boeke and Wang labs for comments and discussion on the work and manuscript. RMM additionally thanks personal support from Xiaoyu Weng.

## Funding

Defense Advanced Research Projects Agency HR0011-17-2-0041 (HHW, JDB)

National Institutes of Health / National Human Genome Research Institute RM1 HG009491 (JDB)

National Science Foundation MCB-1453219 (HHW)

Burroughs Wellcome Fund PATH1016691 (HHW)

Irma T. Hirschl Trust (HHW)

Dean’s Fellowship from the Graduate School of Arts and Sciences of Columbia University (RMM)

## Author contributions

RMM, JT, JDB and HHW developed the initial concept. JT, RMM, AK, SP, HB, SG, LL, MJS, XG, DRJ, and AM performed experiments and analyzed the results. The overall project was supervised by HHW and JDB. The manuscript was drafted by JT, RMM, JDB, and HHW with input from all authors.

## Data and materials availability

Sequencing data generated for this study is deposited in the NCBI SRA at accession number PRJNA742028 (pending).

## EXTENDED DATA

**Figure ED1.**
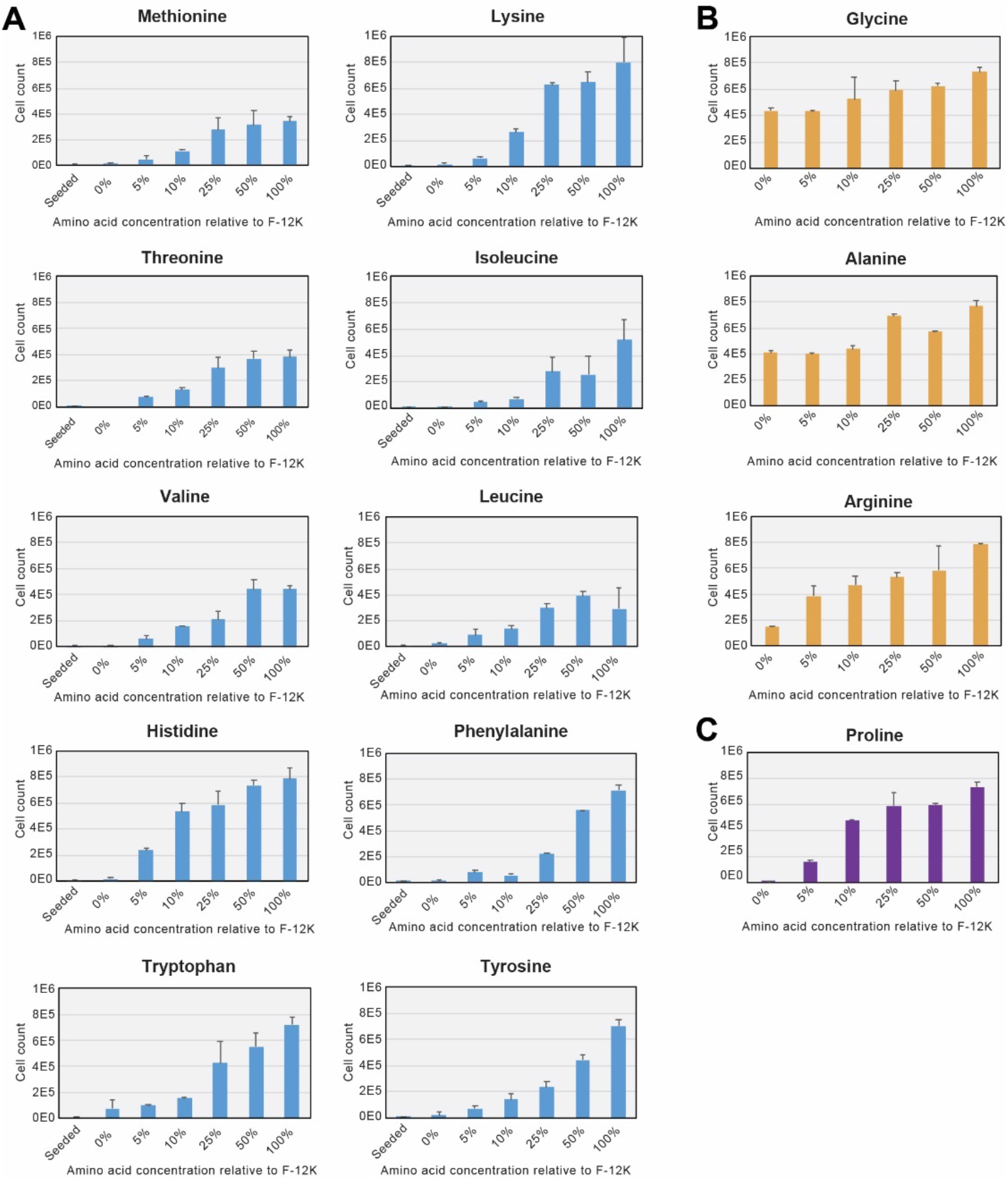
Amino acid dropout growth assays in CHO-K1. **(A)** Growth assays on CHO-K1 cells seeded into media with reduced or omitted essential amino acids. Percentages represent the relative media amino acid concentration compared to standard F-12K medium. 100% corresponds to 0.02 mM (Trp), 0.05 mM (Tyr), 0.06 mM (Ile, Met, Phe), 0.2 mM (Ala, Asn, Asp, Glu, Gly, Leu, Ser, Thr, Val), 0.22 mM (His), 0.4 mM (Cys, Lys), 0.6 mM (Pro), and 2 mM (Arg, Gln), respectively. Error bars show standard deviation of three replicates. **(B)** CHO-K1 cells grow with minimal defects in medium lacking glycine and alanine, but are more sensitive to arginine starvation. **(C)** CHO cells do not grow without exogenous proline, confirming the known epigenetic proline auxotrophy found in this cell line. Error bars show standard deviation of three replicates.

**Figure ED2.**
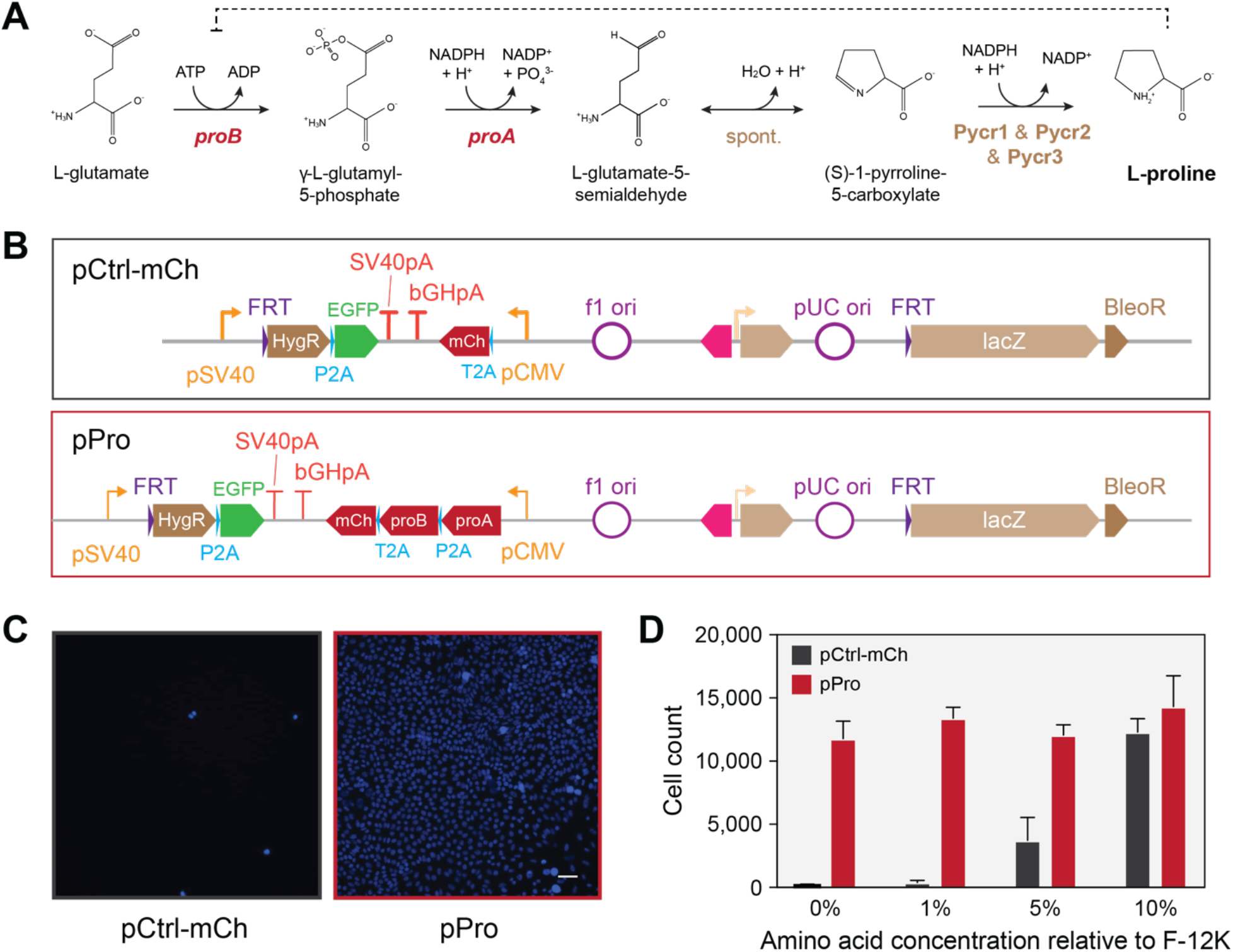
Imported bacterial genes rescue CHO epigenetic proline auxotrophy. **(A)** Proline biosynthetic pathway from glutamate. **(B)** Schematic representation of two constructs to test bacterial gene rescue of of CHO-K1 auxotrophy, one expressing the *E. coli proA* and *proB* genes, and a control construct expressing only mCherry. Constructs are shown as they appear following integration into the FLP-In CHO line. **(C)** Hoechst 33348 live nuclei microscopy shows cells expressing the pPro construct grow in the absence of proline and appear healthy. Scale bar indicates 50 μm and is shared across images **(D)** Growth assay of cells seeded into proline free medium and grown for 5 days. The bacterial gene pPro pathway resulted in robust growth recovery in proline-free medium. Error bars show standard deviation of three replicates.

**Figure ED3.**
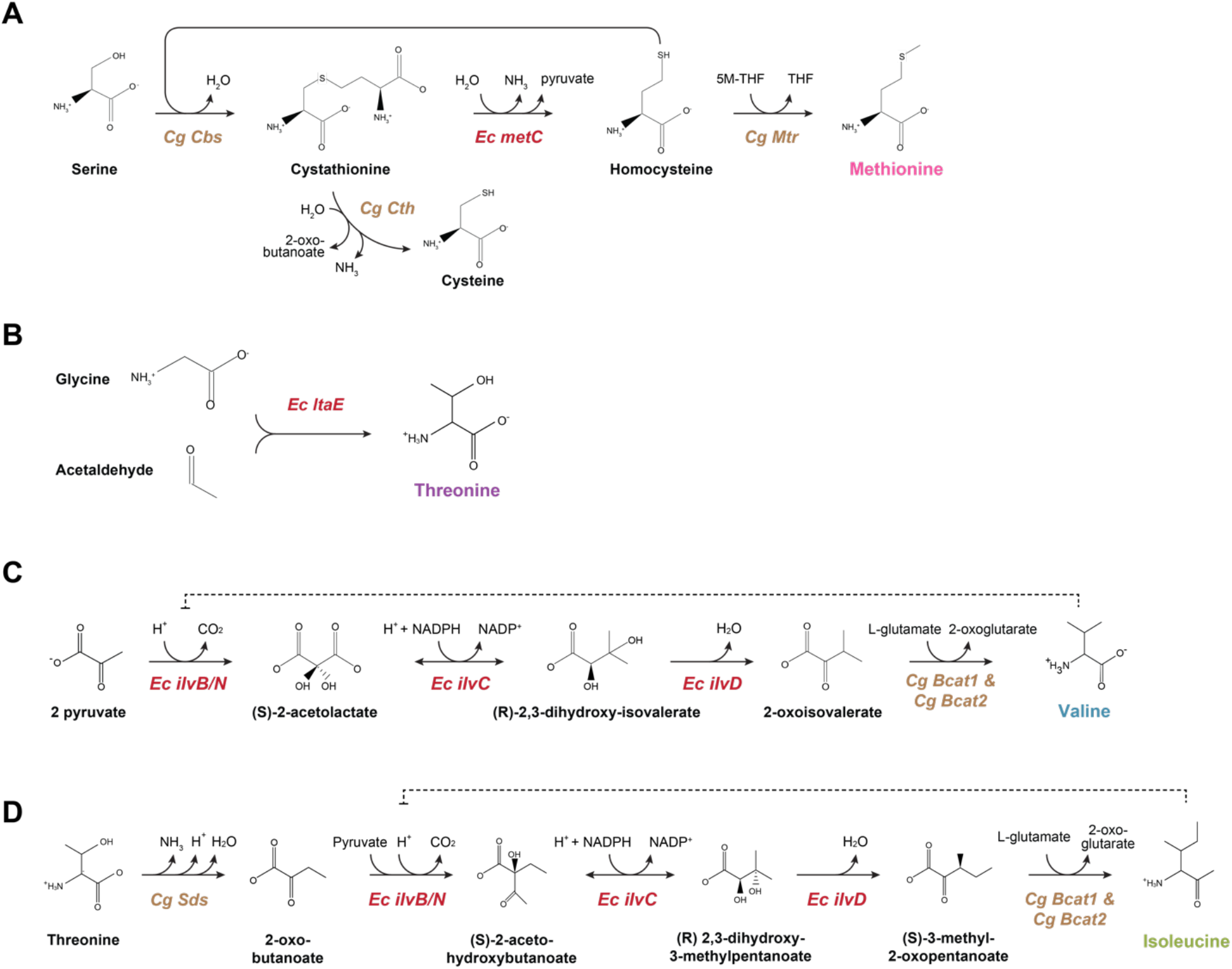
Schematic outlines of the MTIV pathways. **(A)** Pathway for methionine biosynthesis. **(B)** Pathway for threonine biosynthesis. Note the shared multifunctional enzymes in **(C)** and **(D)** responsible for both isoleucine and valine production. In all panels “*Ec*” refers to *E. coli*, and *“Cg”* to *C. griseus*, the native CHO parental organism.

**Figure ED4.**
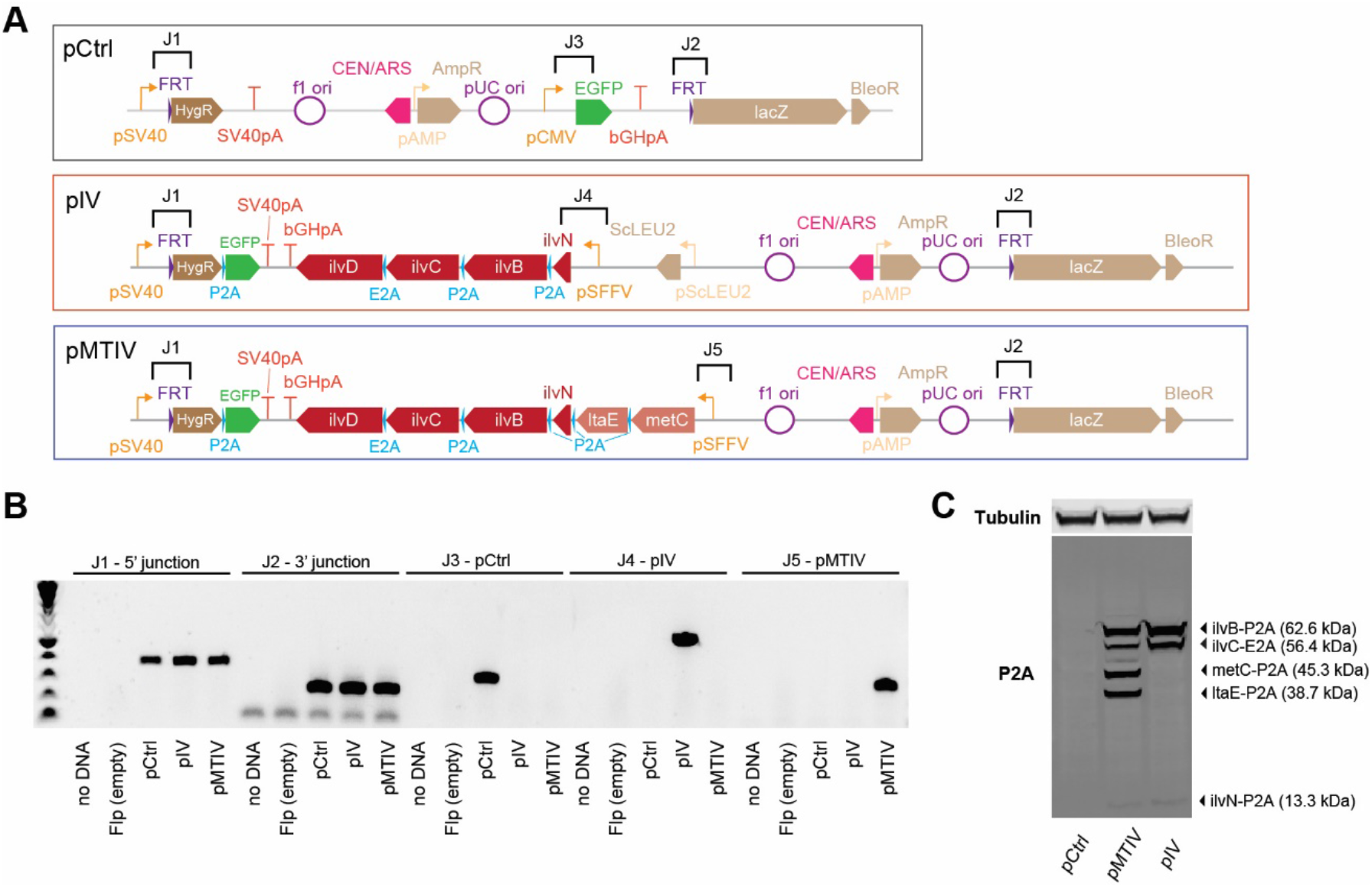
Confirmation of BCAA biosynthetic construct integration and 2A peptide processing by PCR and Western blot. **(A)** Construct were designed with PCR landing pads flanking fragment junctions for rapid screening during construct assembly *in yeasto*, shown above construct diagrams with brackets. **(B)** PCRs across these junctions confirm correct assembly of the control, pMTIV, and pIV constructs. **(C)** Immunoblotting against P2A peptides in pMTIV and pIV cells confirms 2A peptides are processed as expected. Note that *ilvD* (65.5 kDa) is not detectable because it lacks a terminal P2A tag.

**Figure ED5.**
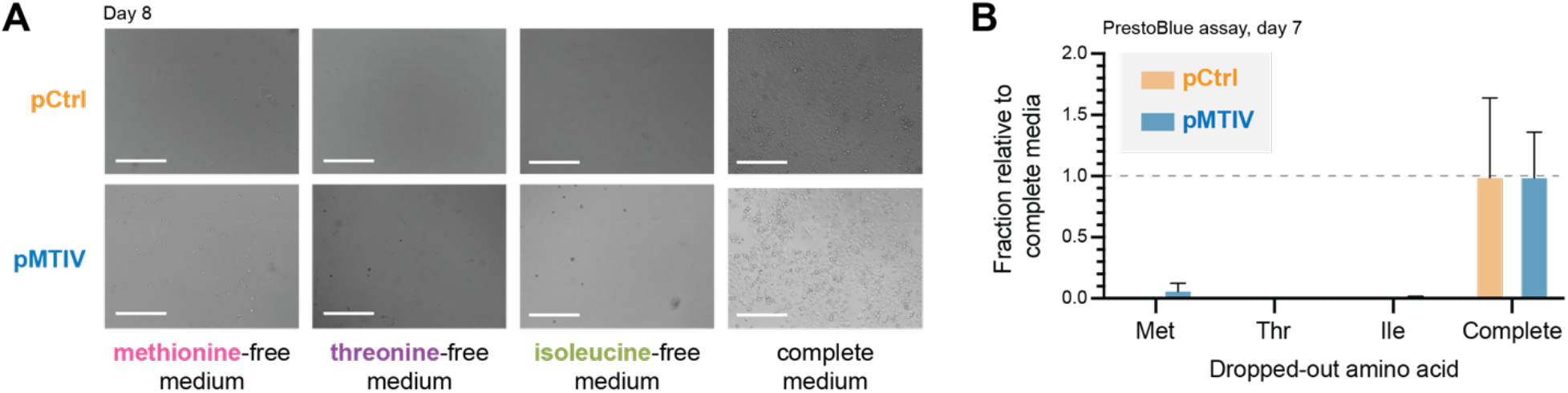
Lack of rescue of methionine, threonine and isoleucine auxotrophy by pMTIV construct in CHO-K1 cells. **(A)** Phase-contrast images of cells carrying either pMTIV and pCtrl constructs after 8 days’ growth on EAA-free media (three leftmost panels) or complete medium (rightmost panel). Scale bar represents 300um. **(B**) PrestoBlue™ cell viability assay of growth on EAA-free or complete media after 7 days’ growth. Error bars show standard deviation of three replicates.

**Figure ED6.**
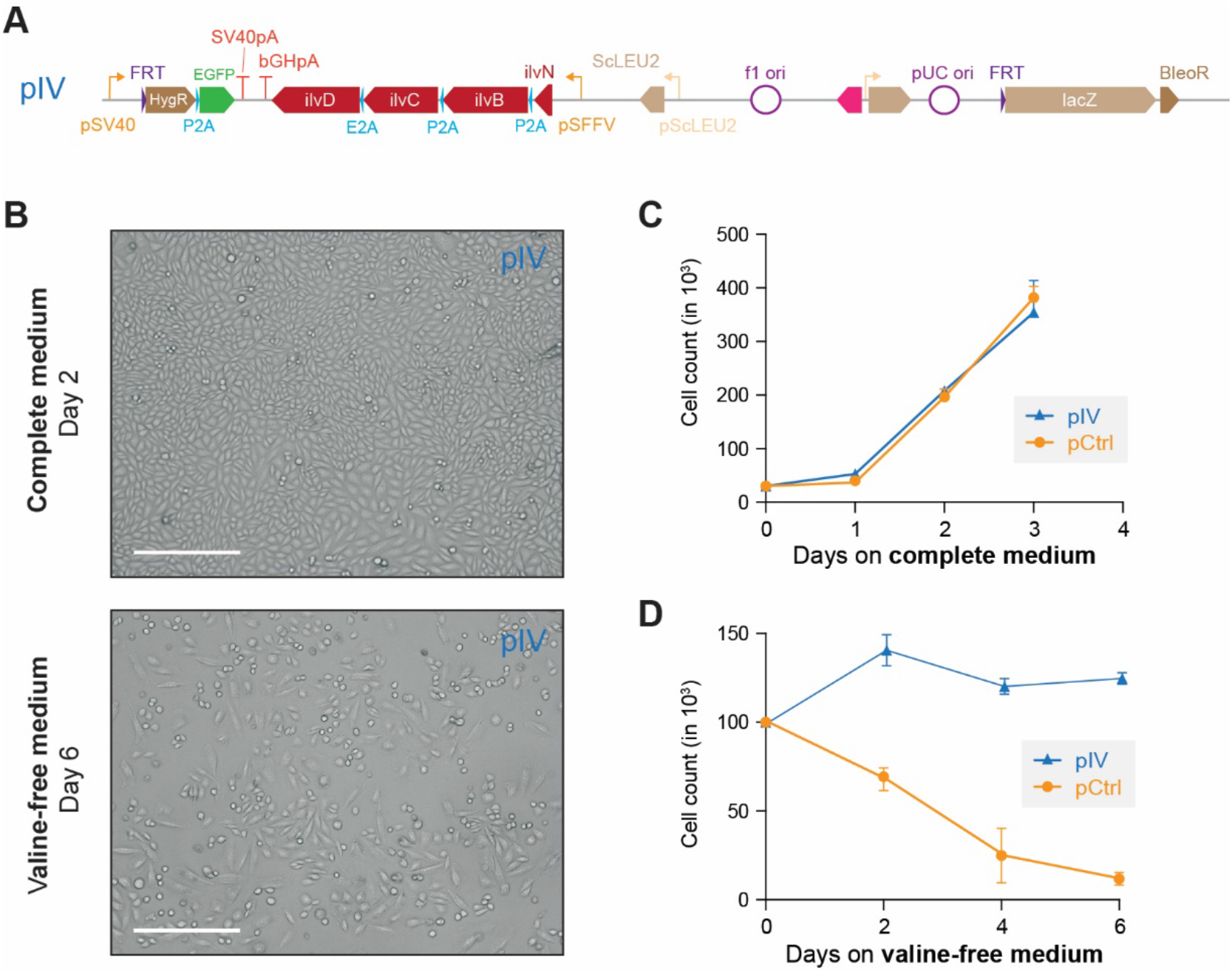
The pIV construct rescues valine auxotrophy. **(A)** The minimized four-gene construct pIV lacking *ltaE* and *metC* after integration into the CHO genome. **(B)** Microscopy of pIV cells in both complete and valine-free medium. The pIV constructs rescue cellular morphology in valine-free medium. Scale bar = 300 um. **(C)** Growth of pIV construct in complete medium. Error bars show standard deviation of three replicates. **(D)** Live cell counting of pIV cells grown on conditioned valine-free medium. Error bars show standard deviation of three replicates.

**Figure ED7.**
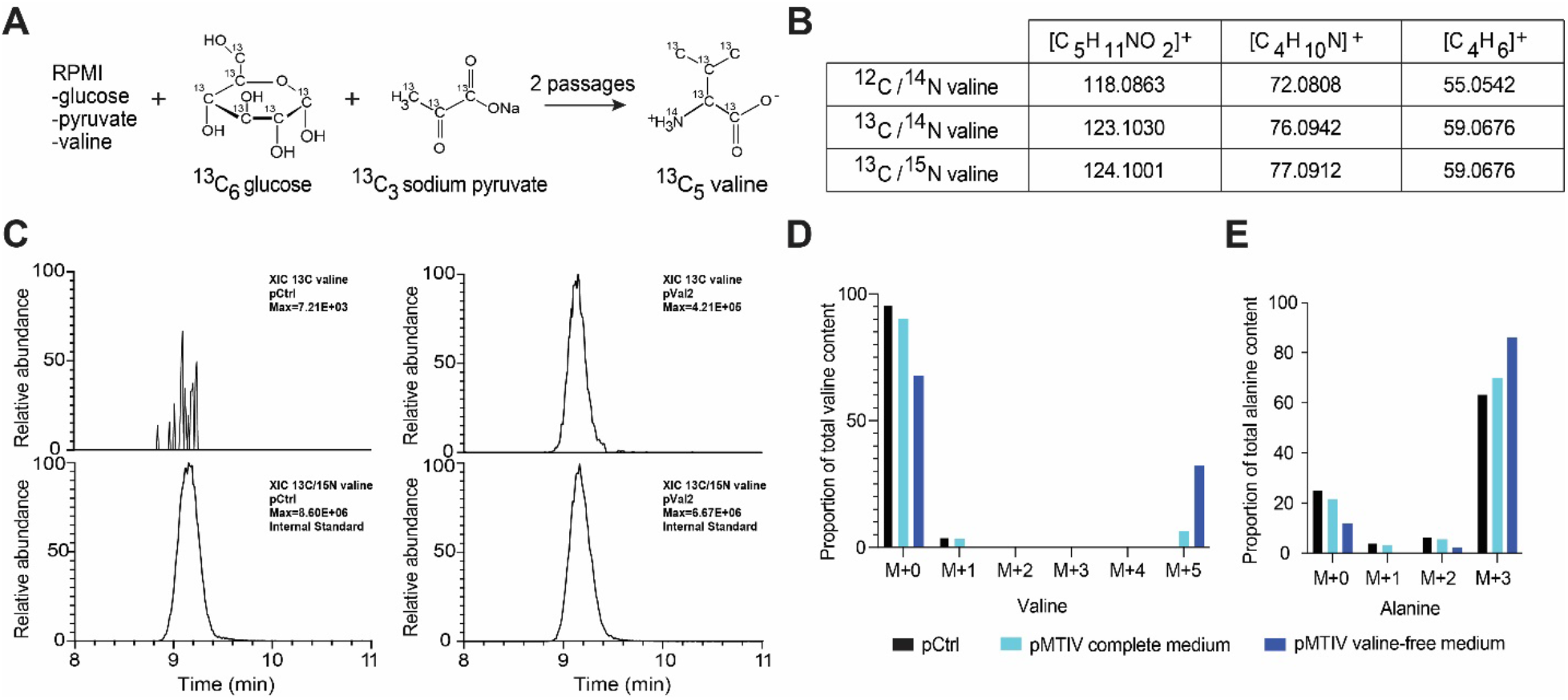
Heavy-labeled metabolomics confirms endogenous valine production. **(A)** Schematic outline of the ^13^C labeling strategy for detection of biosynthesized valine. **(B)** Expected *m*/*z* ratios for ^12^C-valine, ^13^C-valine, and spiked-in internal standard ^13^C/^15^N-valine in unfragmented and fragmented states. **(C)** Extracted ion chromatography shows that the presumed ^13^C-valine found in pMTIV cells runs similarly to spiked-in internal standard ^13^C/^15^N-valine, further confirming that ^13^C-valine is produced in pMTIV cells. The presence of ^13^C-valine is specific to pMTIV cells and absent from control cells **(D)**. In theory, partial ^13^C labeling of valine is possible if just one of the two pyruvate substrates is ^13^C-labelled. Partially ^13^C-labeled (^13^C_2_, ^13^C_3_, ^13^C_4_) valine species were below the level of detection. Therefore, the proportion of ^13^C_5_-valine relative to total valine abundance within the cell can serve as a proxy for the degree to which intracellular valine biosynthesis is meeting cellular valine demands. The lesser occurrence of ^13^C_1_-valine can be explained by the natural abundance of ^13^C. 32.2% of total valine abundance in pMTIV cells cultured in valine-free medium was ^13^C_5_-labeled whereas the corresponding proportion for pMTIV cells cultured in complete medium was 6.4% **(E)** Non-essential amino acid alanine is biosynthesized from pyruvate and is not found in RPMI medium. Alanine can thus serve a proxy for proteome turnover assuming similar usage rates for alanine and valine. 86.1% of alanine was ^13^C_3_ alanine in pMTIV cells cultured in valine-free RPMI medium.

**Figure ED8.**
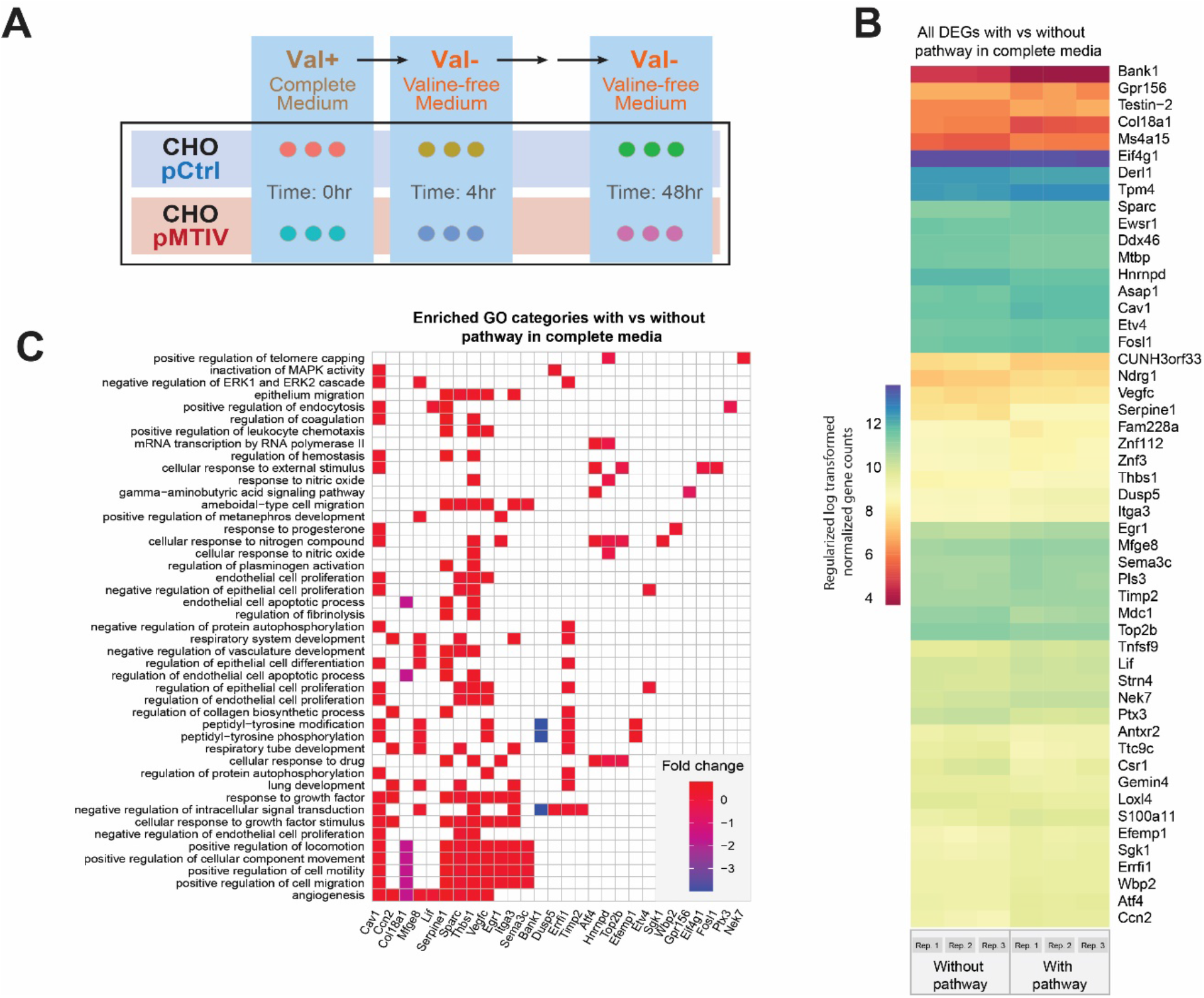
Transcriptomic analysis of pMTIV pathway effect in Complete Medium. **(A)** Schematic representation of the starvation regimen applied prior to collecting samples for transcriptomic analysis. **(B)** Regularized log fold changes in expression across the 51 differentially expressed genes when comparing cells with the pMTIV pathway to those without grown in complete medium. Values from three replicates are plotted side-by-side in each column. **(C)** GO category enrichment among the 51 differentially expressed genes.

**Figure ED9.**
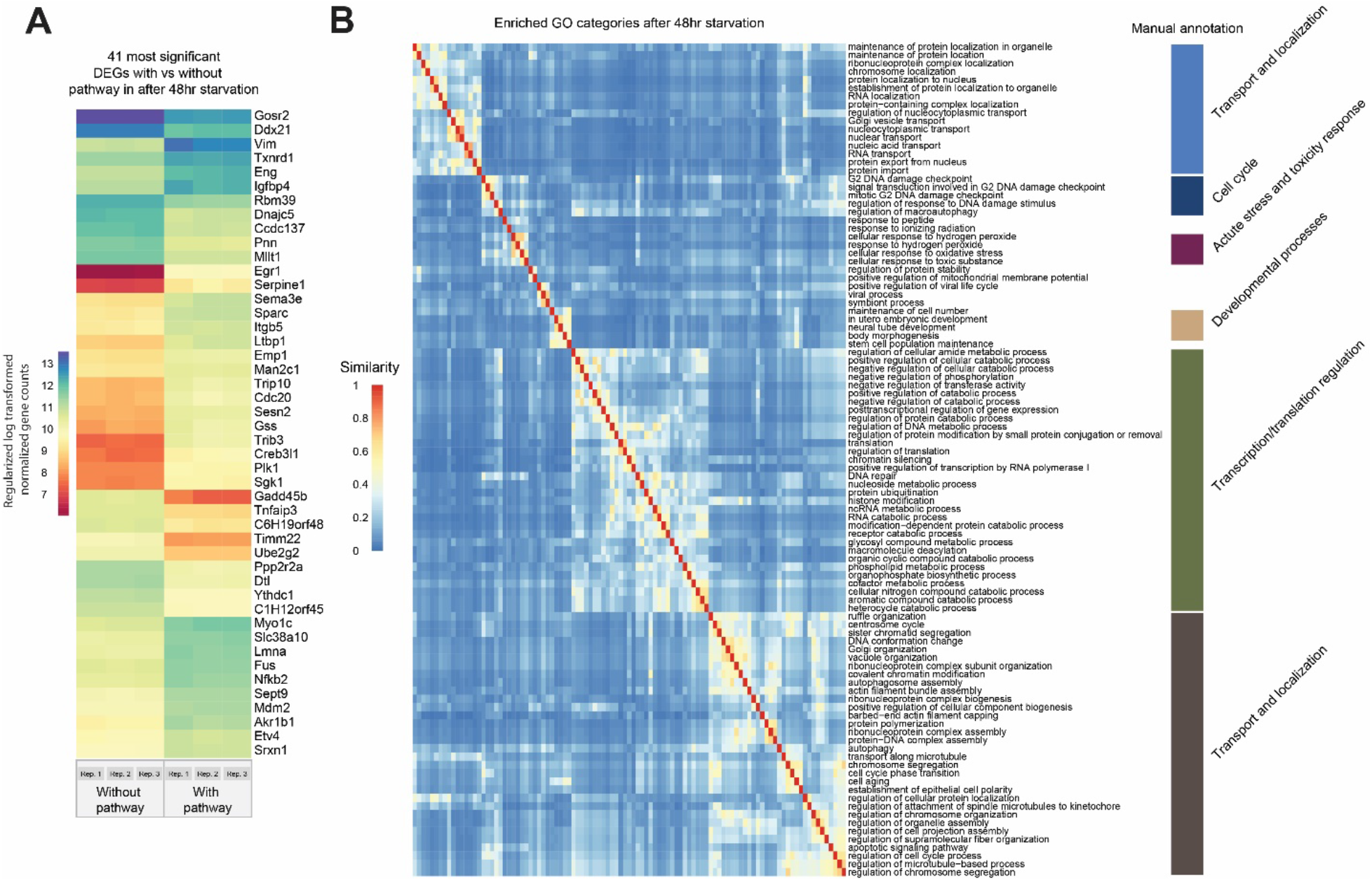
Transcriptomic analysis of pMTIV pathway effect in Valine-free Medium. **(A)** Regularized log transformed expression of the 50 most significant differentially expressed genes when comparing cells with the pMTIV pathway to those without after 48 hours of starvation in valine-free medium. Values from three replicates are plotted side-by-side in each column. **(B)** GO category enrichment among these genes, clustered and colored using a GO ontology distance metric, and manually grouped into larger supercategories, on the right.

**Table ED1.**
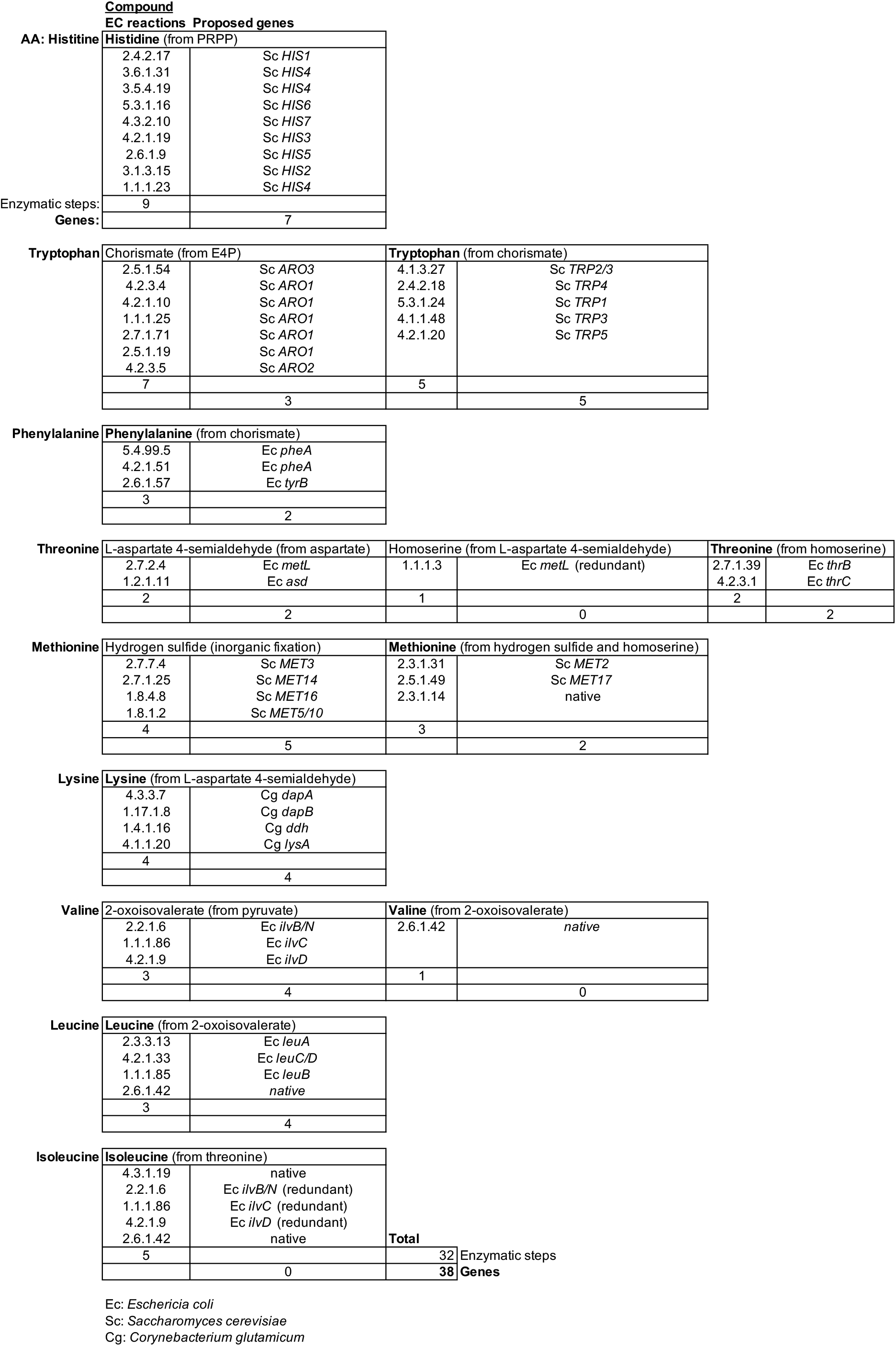
Minimal prototrophic pathways set for AA prototrophy in mammalian cells.

**Table ED2.**
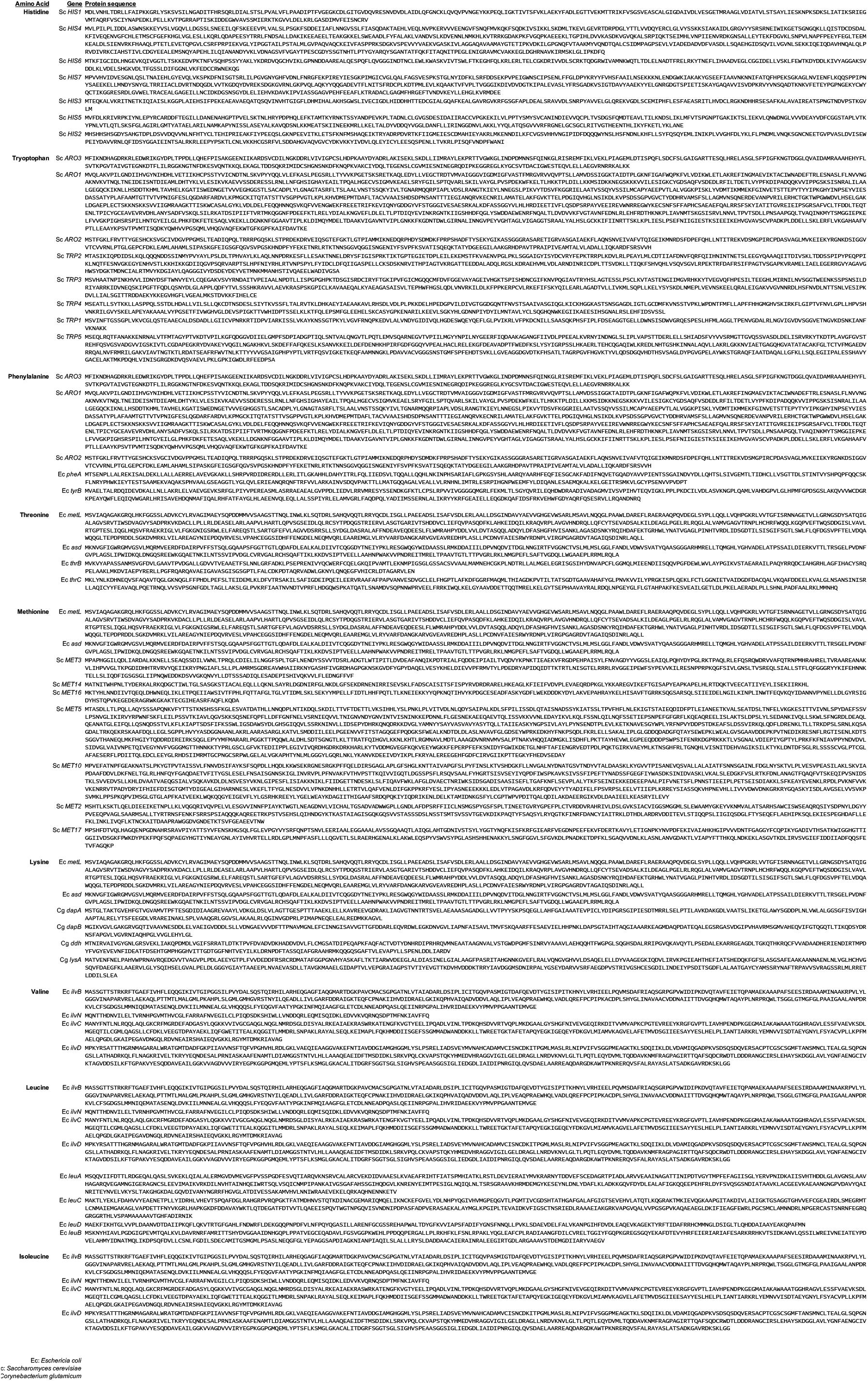
Protein sequence for each gene listed in Table ED1.

**Table ED3.**
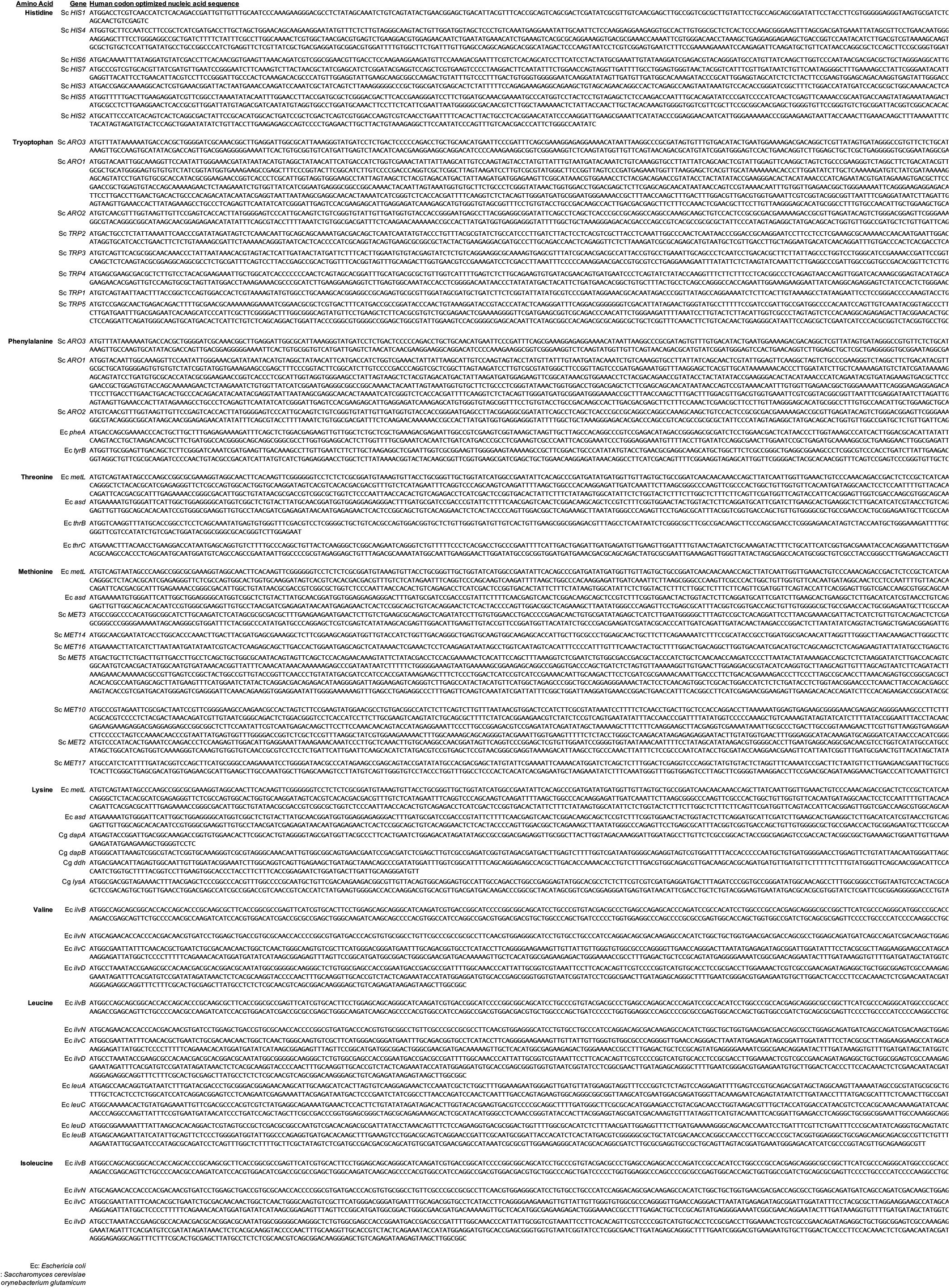
Codon-optimized nucleotide sequence for each gene listed in Table ED1.

**Table ED4.**
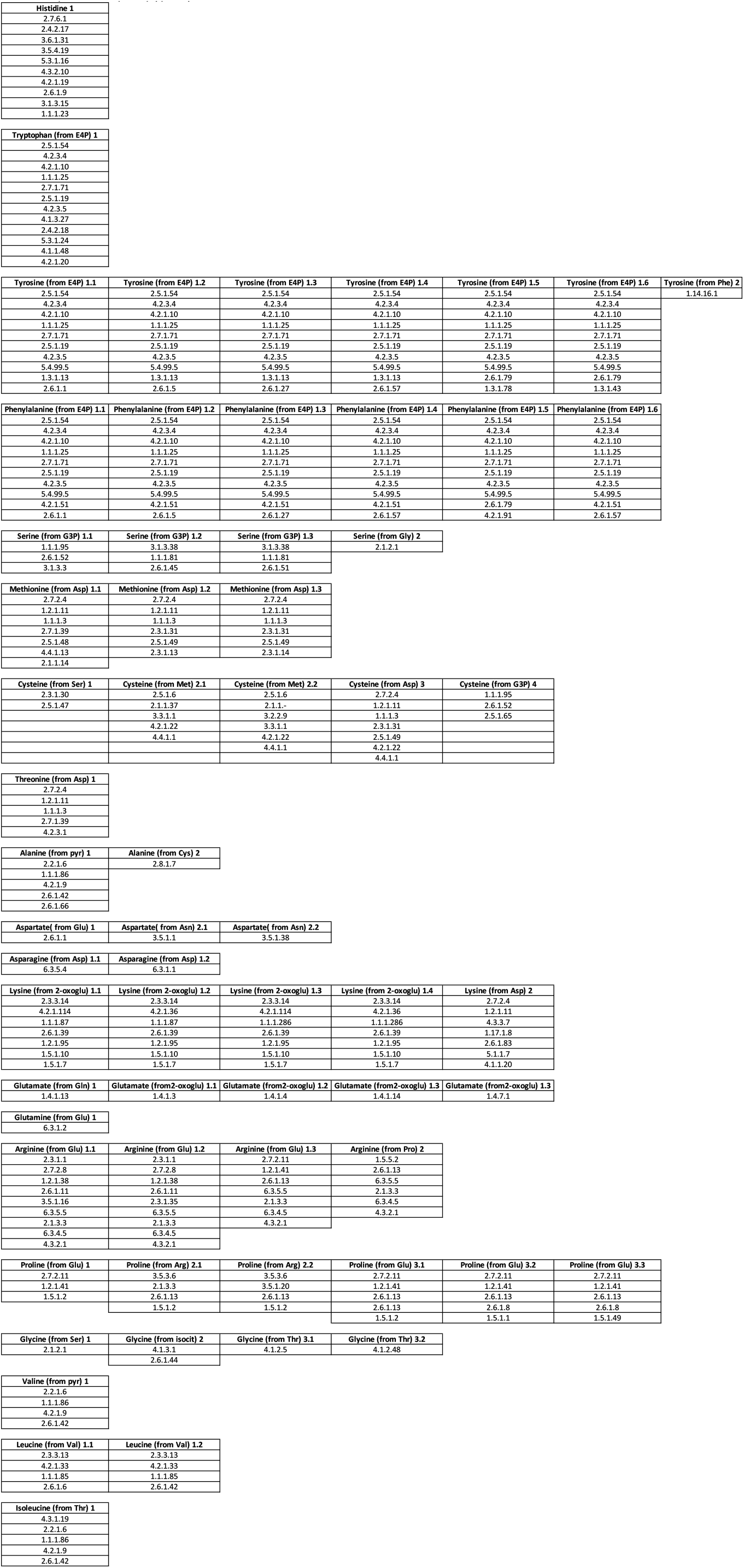
Complete amino acid prototrophy pathway EC numbers Histidine 1.

## REFERENCES

1 Payne, S. H. & Loomis, W. F. Retention and loss of amino acid biosynthetic pathways based on analysis of whole-genome sequences. Eukaryot Cell 5, 272–276, doi:10.1128/EC.5.2.272-276.2006 (2006).

2 Akashi, H. & Gojobori, T. Metabolic efficiency and amino acid composition in the proteomes of Escherichia coli and Bacillus subtilis. Proc Natl Acad Sci U S A 99, 3695–3700, doi:10.1073/pnas.062526999 (2002).

3 Seligmann, H. Cost-minimization of amino acid usage. J Mol Evol 56, 151–161, doi:10.1007/s00239-002-2388-z (2003).

4 Guedes, R. L. et al. Amino acids biosynthesis and nitrogen assimilation pathways: a great genomic deletion during eukaryotes evolution. BMC Genomics 12 Suppl 4, pS2, doi:10.1186/1471-2164-12-S4-S2 (2011).

5 Swire, J. Selection on synthesis cost affects interprotein amino acid usage in all three domains of life. J Mol Evol 64, 558–571, doi:10.1007/s00239-006-0206-8 (2007).

6 Zengler, K. & Zaramela, L. S. The social network of microorganisms - how auxotrophies shape complex communities. Nat Rev Microbiol 16, 383–390, doi:10.1038/s41579-018-0004-5 (2018).

7 Wilson, A. C. & Duncan, R. P. Signatures of host/symbiont genome coevolution in insect nutritional endosymbioses. Proc Natl Acad Sci U S A 112, 10255–10261, doi:10.1073/pnas.1423305112 (2015).

8 Isaacs, F. J. et al. Precise manipulation of chromosomes in vivo enables genome-wide codon replacement. Science 333, 348–353, doi:10.1126/science.1205822 (2011).

9 Mitchell, L. A. et al. Synthesis, debugging, and effects of synthetic chromosome consolidation: synVI and beyond. Science 355, doi:10.1126/science.aaf4831 (2017).

10 Fredens, J. et al. Total synthesis of Escherichia coli with a recoded genome. Nature 569, 514–518, doi:10.1038/s41586-019-1192-5 (2019).

11 Boeke, J. D. et al. GENOME ENGINEERING. The Genome Project-Write. Science 353, 126–127, doi:10.1126/science.aaf6850 (2016).

12 Heng, B. C., Aubel, D. & Fussenegger, M. Prosthetic gene networks as an alternative to standard pharmacotherapies for metabolic disorders. Curr Opin Biotechnol 35, 37–45, doi:10.1016/j.copbio.2015.01.010 (2015).

13 Tan, X., Letendre, J. H., Collins, J. J. & Wong, W. W. Synthetic biology in the clinic: engineering vaccines, diagnostics, and therapeutics. Cell 184, 881–898, doi:10.1016/j.cell.2021.01.017 (2021).

14 Kitada, T., DiAndreth, B., Teague, B. & Weiss, R. Programming gene and engineered-cell therapies with synthetic biology. Science 359, doi:10.1126/science.aad1067 (2018).

15 Fischer, S., Handrick, R. & Otte, K. The art of CHO cell engineering: A comprehensive retrospect and future perspectives. Biotechnol Adv 33, 1878–1896, doi:10.1016/j.biotechadv.2015.10.015 (2015).

16 Szymczak-Workman, A. L., Vignali, K. M. & Vignali, D. A. Design and construction of 2A peptide-linked multicistronic vectors. Cold Spring Harb Protoc 2012, 199–204, doi:10.1101/pdb.ip067876 (2012).

17 O’Gorman, S., Fox, D. T. & Wahl, G. M. Recombinase-mediated gene activation and site-specific integration in mammalian cells. Science 251, 1351–1355, doi:10.1126/science.1900642 (1991).

18 Hefzi, H. et al. A Consensus Genome-scale Reconstruction of Chinese Hamster Ovary Cell Metabolism. Cell Syst 3, 434–443 e438, doi:10.1016/j.cels.2016.10.020 (2016).

19 Cunningham, J. A., Liu, A. G., Bengtson, S. & Donoghue, P. C. The origin of animals: Can molecular clocks and the fossil record be reconciled? Bioessays 39, 1–12, doi:10.1002/bies.201600120 (2017).

20 Amorim Franco, T. M. & Blanchard, J. S. Bacterial Branched-Chain Amino Acid Biosynthesis: Structures, Mechanisms, and Drugability. Biochemistry 56, 5849–5865, doi:10.1021/acs.biochem.7b00849 (2017).

21 Zhenyukh, O. et al. High concentration of branched-chain amino acids promotes oxidative stress, inflammation and migration of human peripheral blood mononuclear cells via mTORC1 activation. Free Radic Biol Med 104, 165–177, doi:10.1016/j.freeradbiomed.2017.01.009 (2017).

22 Zhang, Y. et al. Expression of threonine-biosynthetic genes in mammalian cells and transgenic mice. Amino Acids 46, 2177–2188, doi:10.1007/s00726-014-1769-0 (2014).

23 Rees, W. D. & Hay, S. M. The biosynthesis of threonine by mammalian cells: expression of a complete bacterial biosynthetic pathway in an animal cell. Biochem J 309 (Pt 3), 999–1007, doi:10.1042/bj3090999 (1995).

24 Black, J. B., Perez-Pinera, P. & Gersbach, C. A. Mammalian Synthetic Biology: Engineering Biological Systems. Annu Rev Biomed Eng 19, 249–277, doi:10.1146/annurev-bioeng-071516-044649 (2017).

25 Gilbert, L. A. et al. Genome-Scale CRISPR-Mediated Control of Gene Repression and Activation. Cell 159, 647–661, doi:10.1016/j.cell.2014.09.029 (2014).

26 Weingarten-Gabbay, S. et al. Systematic interrogation of human promoters. Genome Res 29, 171–183, doi:10.1101/gr.236075.118 (2019).

27 Lim, A. R., Rathmell, W. K. & Rathmell, J. C. The tumor microenvironment as a metabolic barrier to effector T cells and immunotherapy. Elife 9, doi:10.7554/eLife.55185 (2020).

28 Tan, C. S. et al. Positive selection of tyrosine loss in metazoan evolution. Science 325, 1686–1688, doi:10.1126/science.1174301 (2009).

29 Dancis, J., Levitz, M. & Westall, R. G. Maple syrup urine disease: branched-chain keto-aciduria. Pediatrics 25, 72–79 (1960).

30 Blau, N., van Spronsen, F. J. & Levy, H. L. Phenylketonuria. Lancet 376, 1417–1427, doi:10.1016/S0140-6736(10)60961-0 (2010).

31 Kemmer, C. et al. Self-sufficient control of urate homeostasis in mice by a synthetic circuit. Nat Biotechnol 28, 355–360, doi:10.1038/nbt.1617 (2010).

32 Ye, H. et al. Pharmaceutically controlled designer circuit for the treatment of the metabolic syndrome. Proc Natl Acad Sci U S A 110, 141–146, doi:10.1073/pnas.1216801110 (2013).

33 Pacold, M. E. et al. A PHGDH inhibitor reveals coordination of serine synthesis and one-carbon unit fate. Nat Chem Biol 12, 452–458, doi:10.1038/nchembio.2070 (2016).

